# Best-of-*n* decision making by human groups

**DOI:** 10.1101/2025.07.23.666271

**Authors:** Nicolas Coucke, Marco Dorigo, Axel Cleeremans, Mary Katherine Heinrich

## Abstract

Collective decision making is a fundamental aspect of group behavior in both animals and humans, and often involves reaching a consensus on the best of *n* options, using empirical evidence. Although many parallels have been drawn between human and animal collective decisions, collective human behavior is rarely studied in the type of embodied scenarios that animals are often faced with. In this study, we placed human groups in a virtual setup similar to nest site selection in social animals, in which they explored a shared environment and reached a consensus based on their observations of empirical features. In groups of up to 10, participants had to reach consensus on the empirically largest of four candidate sites without verbal communication, instead using movement-based interactions in a custom-developed 3D virtual environment for online multi-participant experiments. The results showed that the speed and accuracy of consensus was importantly modulated by perceptual difficulty and information availability, but that no speed–accuracy trade-off was present. Participants attempted to reach consensus on the empirically largest site by flexibly adapting their use of social information to perceptual difficulty, their spatial position, and the time already spent supporting some option. When a minority of informed individuals were present, these individuals exercised greater independence and influenced the group to faster and more accurate consensus. These results extend previous findings on social decision making strategies in humans to nonverbal scenarios akin to those of social insects.

## Introduction

The ability to perform cohesive collective actions by reaching a consensus is crucial to the survival of many social animal species [1]. Ants, bees, birds, and primates all integrate separate observations made by many individuals in the process of coming to a consensus, for example on nest-site selection or when and where to travel next [2–5]. Similarly, in human groups, coherent collaborative actions require the integration of different viewpoints through collective decision-making processes [6].

Some characteristics of social animal collective decision making have been compared to human collective decision making [7, 8], and it has even been suggested that social animals could inspire the development of new decision-making frameworks in humans, especially in the context of democratic processes [7, 9]. Recently, it has also been shown that simple social animals have stronger collective benefits than humans during a similar task: ants performed an object transport task more effectively when working collectively, while human groups showed almost no benefit [10]. However, because humans typically use verbal communication, human and animal best-of-*n* decision-making processes have usually been studied in very different contexts, complicating the potential transferability of findings and mechanisms between species. In humans, best-of-*n* decision making is typically studied as a verbal exchange of information or arguments, mostly studied for discrete, symbolic opinions [6, 11, 12], while in animals it is studied as a series of embodied interactions in a shared physical environment.

A notable exception is collective motion: studies of human crowd dynamics [13–17] resemble studies of social animals in which groups remain together while moving according to nonuniform goals [18–20]. In this type of study, multiple subgroups are given different movement goals, which individuals need to balance with the goal of staying close to the group in order to move cohesively. In both the human and animal results, the groups reached a consensus more often when a proportion of the group was initially unopinionated.

In site selection, reaching a best of-*n* consensus requires arbitrating different movement goals while also exploring and gathering information from the environment. For example, when ants or honeybees select a new nest site, individuals explore possible sites in the environment and interact with one another to pool the information they obtained [2, 3, 21, 22]. The balance of information gathering (exploring sites) and information pooling (interacting with one another in order to reach a consensus) is important to success, and the two processes are often balanced actively (e.g., in ants and honeybees [21, 23]). Although human groups often coordinate to make much more complex decisions than those made by social insects, their processes still generally involve information gathering and information pooling [24].

In this paper, we study how human groups balance the gathering of information in an environment with pooling it to reach a consensus. We study this question using a site selection scenario that resembles the settings to which social animals are usually exposed while foraging or during nest selection. In our experimental setup, participants need to reach a consensus to select the largest of four candidate sites, while not having any communication channel other than observing each others’ positions. They need to explore their shared environment to gather information and can indicate their choice by positioning themselves at a certain site. In order to succeed at the task, all group members must simultaneously position themselves at the correct site. In this setup, if all participants could easily perceive that one option is superior (one site is clearly far larger than all others), consensus could be reached using only individual observations. However, when perceiving the empirical difference between sites is difficult, individuals will likely have to form a consensus despite initial disagreement. They will thus have to take into account social information to some degree in their decision making.

Humans (and other animals) make use of several social learning strategies to regulate their use of social and individual information [25]. For example, when making decisions based on empirical observations, individuals have been shown to copy the opinions of peers more often when the task becomes more difficult [26]. Similarly, when seeking rewards in a shared environment, individuals have been shown to make a trade-off between individual exploration and copying their peers when tasked with maximizing their individual rewards [27, 28]. In this paper, we investigate how social and individual information are regulated when participants are required to reach a consensus. For example, when an individual in our site selection setup has not yet made enough direct observations to sufficiently inform an opinion, or simply feels unsure about those observations, it might be judged more effective to follow the current majority opinion than to explore further.

To select a site, an individual must move around in the environment. These movements might importantly modulate social information use, for example when an individual has already spent time moving towards one option. As previously found for individual decision making [29, 30], an individual might be less likely to change opinion (and thus movement direction) when already near a selected site. Similarly, due to a sensitivity to sunk costs [31], an individual might be reluctant to change opinion after already spending a long time located at a certain site.

Humans also use social learning strategies to select which individuals to copy or follow, for example due to the individual’s degree of past accuracy or displayed level of confidence [32]. In the setting of individual reward gathering in a shared environment, participants inferred that peers were confident in the reward quality of a certain location when they lingered there [28]. In a setting that required a group to come to a consensus on a site but did not require exploration of the environment beforehand, confidence was signaled by moving to a site quickly and remaining there persistently, even despite a disagreeing majority [33]. Similarly, in human crowd dynamics, more informed individuals gained social influence by moving early and consistently [15, 16]. In the present study, we investigate how informed individuals exert influence on others during consensus-reaching, based on empirical observations of a shared environment.

## Experiment design

We designed a best-of-*n* collective decision-making task with human groups that control identical avatars in a 3D virtual environment. Each participant controls one avatar that can move omnidirectionally in two dimensions around a virtual environment with planar ground. The participant has a first-person point-of-view from their avatar, in the direction the avatar is moving, and can observe both the other avatars and their shared environment (see Fig. 1). Participants did not have any communication channel apart from observing the two-dimensional movements of each other’s avatars.

**Fig 1.**
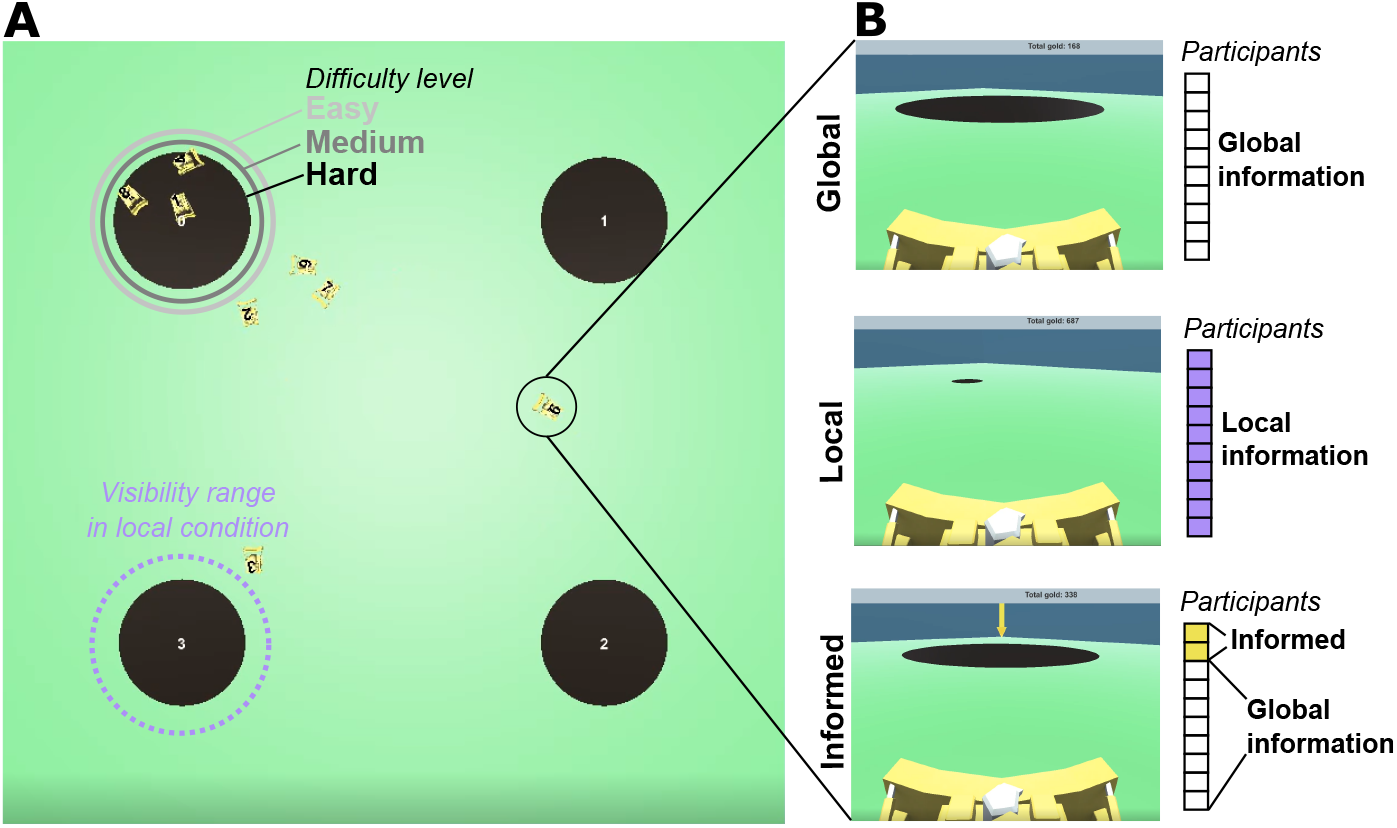
Experiment design. Participants control robot avatars (small yellow bulldozers) and can reach a correct consensus by placing all their avatars together on the largest of four circular sites in the environment (dark brown circles). **A) Zenithal view (only visible to the experimenter) of an example *hard* trial**. The size difference between correct and incorrect sites is largest in the *easy* difficulty trials and smallest in the *hard* difficulty trials (light pink, dark pink, and black lines around top-left site). In the *local* condition, participants can only see the site sizes when approaching them closely (light purple dotted line around bottom-left site). **B) Images show example first-person views that participants can see from their avatars, in the three different experiment conditions. The adjacent cells represent the information available to participants in the group**. In the *global* condition, all participants can see the sites locations and sizes from a distance (top frame). In the *local* condition, all participants can see only the sites locations (and not their sizes) from a distance (middle frame); they must approach a site closely to see its size (the range is labeled in A). In the *informed minority* condition, all participants can see the site locations and sizes from a distance, like in the *global* condition, but two of the participants are informed of the correct (i.e., largest) site by a yellow arrow above it (bottom frame).

Our task is inspired by the best-of-*n* decision-making scenarios often studied in natural and artificial swarm intelligence, in which agents have to collectively select one of *n* options in the environment [34]. In social insects, best-of-*n* decision making has been primarily studied in the context of nest-site selection in a few species of honey bees and ants. These species generally select nest sites via the same basic process [35]: some proportion of the colony acts as scouts, which individually explore and discover potential sites, then pool information about their quality assessments until a quorum-sensing mechanism triggers the movement of the colony to a new site. In natural settings, the potential sites are numerous (e.g., in honeybees, the potential sites can consist of many trees located in a large open area or several regions within a larger area [36, 37]). However, in experimental settings, the available sites have usually been restricted to the range of *n* = {2, …, 7} to investigate the constituent mechanisms in a more detailed manner—e.g., to study the trade-off between site quality and distance [38] or different aspects of quality such as humidity and light [39], the probability of choosing the best site depending on its quality and the qualities of other sites [40, 41], the individuality versus collectivity of site comparison mechanisms [42], and symmetry breaking when sites are of equal or similar quality [43]. Likewise, the vast majority of best-of-*n* decision-making studies in swarm robotics have also used the range of *n* = {2, …, 7}, for example to study the effects of dynamic site qualities [44], site layout [45], malicious or stubborn robots [46, 47], miscommunication [48], and negative updating [49]. In general, the greater the number of sites to choose among, the greater the task difficulty [43, 50]. However, only *n* = 2 has been shown to have qualitatively different decision-making dynamics than other *n* sizes [43]; when studied, dynamics for any *n >* 2 have been shown to be consistent in their macrophases [43]. In this paper, we have selected *n* = 4 to represent the *n >* 2 category; *n* = 4 should represent a near-median task difficulty for the typically studied range of *n* = {2, …, 7} .

In our setup, there are *n* = 4 potential sites, with one site being of higher quality and *n* − 1 sites being of lower quality. In our setup, groups of 7–10 participants are tasked with collectively selecting the largest of four possible candidate sites, within a time limit of 100 seconds. Participants start with their avatars near the center of the environment, with the four candidate sites equidistant from the center. Participants can reach a consensus by all group members placing their avatars on top of one of the four candidate sites. Participants are rewarded when they reach a correct consensus, that is, a consensus on the largest site. Also, the faster they reach a correct consensus, the greater their reward. If they reach an incorrect consensus or time runs out, they receive no reward.

We implemented several experimental manipulations. To explore how people balance individual and social information, we first used trials with three different difficulty levels (*easy, medium*, and *hard*), which corresponded to the empirical size differences between the correct and incorrect candidate sites (radius of the correct site is 8.3%, 16.6%, or 25% larger than the incorrect sites, respectively). Secondly, we manipulated the ease of accessing empirical information, with a *global* condition in which the size of the sites could be observed from far away and a *local* condition in which a site had to be approached closely to observe its size (see Fig. 1). Then, to investigate the influence of informed individuals on consensus, we added an *informed minority* manipulation of the *global* condition, in which two individuals (in groups of 7–10) were informed of the correct site at the beginning of the trial, by an arrow displayed above the correct site. The informed individuals remained indistinguishable from the other avatars: uninformed individuals did not know whom was informed nor in which trials informed individuals were present.

## Results

In total, we conducted 554 trials across 32 experimental sessions, with group sizes between 7 and 10 participants. A consensus decision was reached within the 100 s time limit in 469 trials and that consensus was correct (successful) in 412 trials (see S1 Appendix for details). Qualitative differences in participant movements between trajectories can be observed in Fig. 2D-F. Qualitative differences between experimental manipulations were evaluated using either linear mixed models (LMMs) or generalized linear mixed models (GLMMs) with a binomial response variable (see Materials and Methods for details). Time series of individuals’ opinions were quantified based on their positions and movements (see Materials and Methods for details).

**Fig 2.**
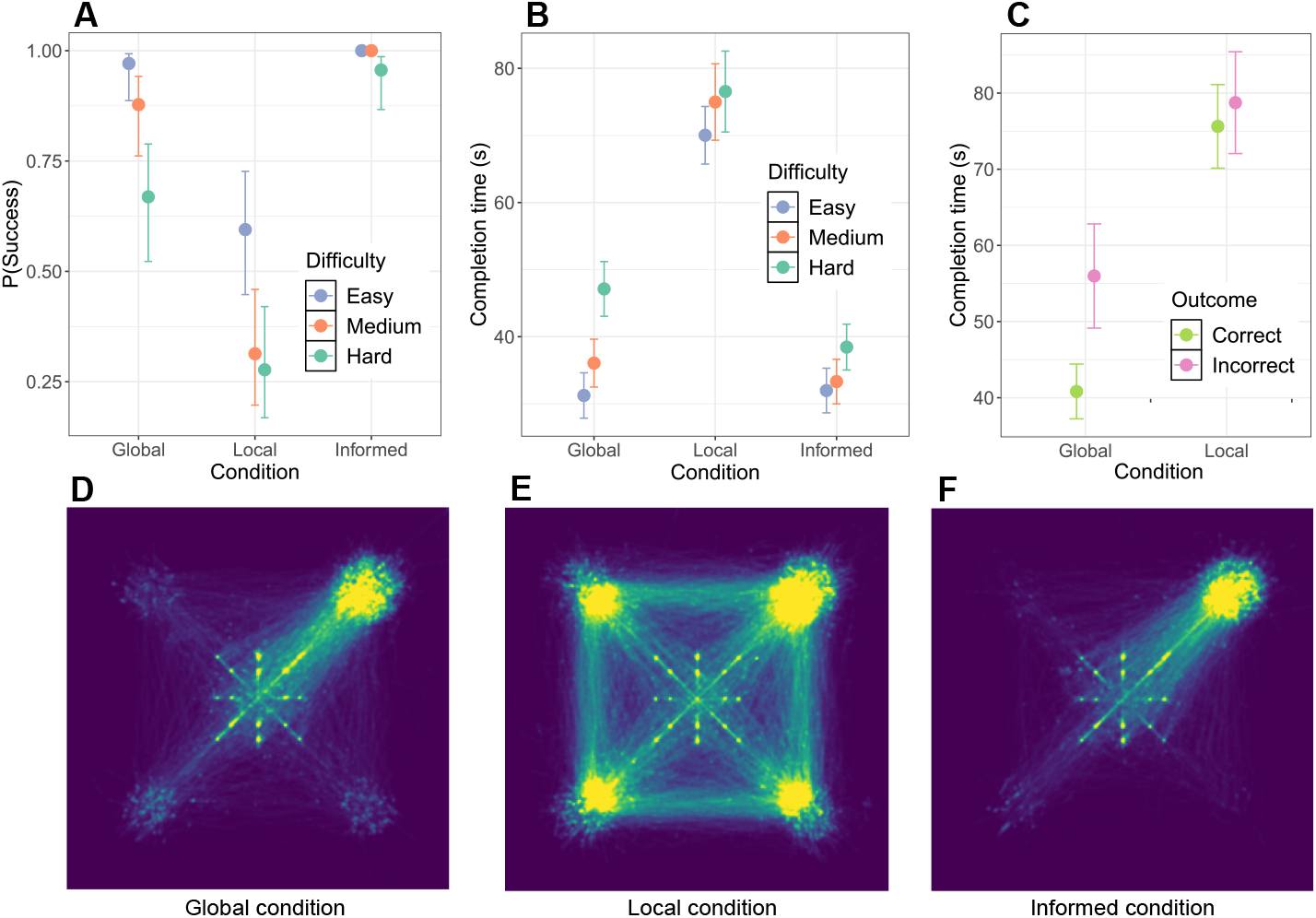
Likelihood and speed of reaching a correct consensus. A) The probability of success (i.e., reaching a correct consensus within the time limit) for each experimental condition and difficulty level, based on all 554 trials (mean and 95% error bar). In all conditions, the groups on average performed better than chance (i.e., *>*25% success rate, as there are 4 options). B) The estimated time to reach a correct consensus in successful trials, for each experimental condition and difficulty level (mean and 95% error bar). C) The estimated time to reach a consensus in *hard* difficulty trials, in both correct (successful) and incorrect trials. Shorter time to reach a consensus for correct trials suggests the absence of a speed–accuracy trade-off. The *informed minority* condition is not shown, as there were no trials ending in incorrect consensus. D-F) Heatmaps of experimental conditions, with the correct candidate site in the top right corner of the arena. Each heatmap shows the trajectories of all participants in all trials of the respective condition (brighter areas indicate higher density). Note that the star-shaped configuration of dots in the center are the starting positions of the avatars. D) In the *global* condition, density is highest on and towards the correct candidate site, with some density distributed among the other three. E) In the *local* condition, density is much more distributed across the correct and incorrect sites. F) In the *informed minority* condition, density is more concentrated on the correct site than in any other condition.

In this section, we first report the process of reaching consensus and the factors influencing consensus speed and accuracy. Next, we report individual behaviors: factors influencing changes of mind, behavior when part of an informed minority, and finally intentional persuasion.

### Reaching consensus on the empirically best option

Firstly, as expected, we found that both the accuracy and the speed of group decision making decreased with increased difficulty (Fig. 2A-B). Across experimental conditions, both the probability of success (*χ*^2^(2) = 25.1, *p <* 0.0001) and the time to reach a consensus (*χ*^2^(2) = 40.7, *p <* 0.0001) were modulated by the difficulty of distinguishing between the empirical sizes of the sites. The *hard* difficulty was less likely to result in success than the *easy* difficulty in both the *local* (*z* = − 3.2, *p* = 0.0036) and *global* (*z* = − 3.6, *p* = 0.001) conditions, with a success rate only slightly better than chance (*>*25%) in the *local* condition. Also, the *hard* difficulty trials took longer to reach a consensus than the *easy* difficulty trials in all conditions (all *p* values *<* 0.02).

Also as expected, the results confirmed that limited access to environmental information hindered the decision-making process. Trials in the *local* condition were less likely to be successful than in the *global* condition (*z* = − 7.2, *p <* 0.0001) and, on average, groups took longer to reach a consensus in the *local* condition (72.7 s *±* 12.8 s) than in the *global* condition (36.3 s *±* 14.6 s). Moreover, the effect of difficulty levels on the time to reach a consensus was less pronounced in the *local* condition than in the *global* condition (*β*_local,hard_ = − 9.36, *t* = − 2.08, *p* = 0.038), suggesting that the time to reach a consensus in the *local* condition was largely determined by the time needed to move between options when exploring the environment. This is also apparent from the average distance between participants’ avatars during the whole trial, which was larger in the *local* condition (see S2 Appendix).

In the *informed minority* trials, the informed individuals effectively led the uninformed majority to the correct candidate site. The *informed minority* trials were almost always successful (98 % of trials) and the probability of success was not modulated by the difficulty levels (*z* = −0.2, *p* = 0.9902). In the *informed minority* trials, consensus was also reached faster (*z* = −2.53, *p* = 0.01), and the average distance between avatars was smaller (see S2 Appendix), than in the *global* and *local* trials. Still, in the *informed minority* condition, participants took longer to reach consensus in *hard* trials than in *easy* trials, although the effect of difficulty on consensus time was less strong in the *informed minority* condition than in the standard *global* condition (*β*_informed,hard_ = − 9.41, *t* = − 2.70, *p <* 0.007). This non-zero effect of difficulty suggests that decisions still occurred faster when the uninformed individuals could easily observe differences between options.

Overall, a speed–accuracy trade-off is notably absent in the results (Fig. 2C). We compared the average time to reach a consensus for trials with a correct versus incorrect consensus, considering only trials of the *hard* difficulty (in the *local* and *global* conditions), because these were the only conditions with a large enough number of incorrect consensus trials. Post-hoc group comparisons indicate that correct trials were completed faster than incorrect trials in the *global* condition (*t* = − 2.477, *p* = 0.0139). The *global* condition is an especially useful comparison case because only 3 trials (approx. 5%) ended without a consensus. No difference was found in the *local* condition. See S4 Fig in S3 Appendix for a more detailed summary of the time to reach a consensus of correct and incorrect trials per group.

### Social information, environment, and movement drive changes of mind

Participants often did not end up at the option they initially moved towards, instead switching opinions either while moving towards an option or after they had already arrived at an option but before a consensus was reached. The probability of switching opinions at any point in time during the trial varied across experimental conditions and difficulty levels (see S4 Appendix). We assessed to what degree switches of opinions were based on social information, and whether social information use was modulated by the difficulty of the trial. For this assessment, we used a measure of disagreeing social information (DeltaSocial). This measure quantifies an individual’s exposure to social pressure to go to an alternative option. It is defined as the difference between the maximum number of group members supporting any one alternative option versus the number of individuals agreeing with the individual’s current selection (see Materials and Methods).

In the *global* condition (Fig. 3A), opinion switches during movement became more likely as more individuals supported the respective alternative (*χ*^2^ = 199.34, *p <* 0.0001), confirming that social information was used. The dependence on social information interacted with the difficulty of the trials (*χ*^2^ = 8.74, *p* = 0.01): participants relied more on social information in the higher difficulty level conditions (*β*_medium_ = 0.11507, SD = 0.04, *t* = 2.78, and *β*_hard_ = 0.12, SD = 0.04, *t* = 3.10, both *p* values *<* 0.006).

**Fig 3.**
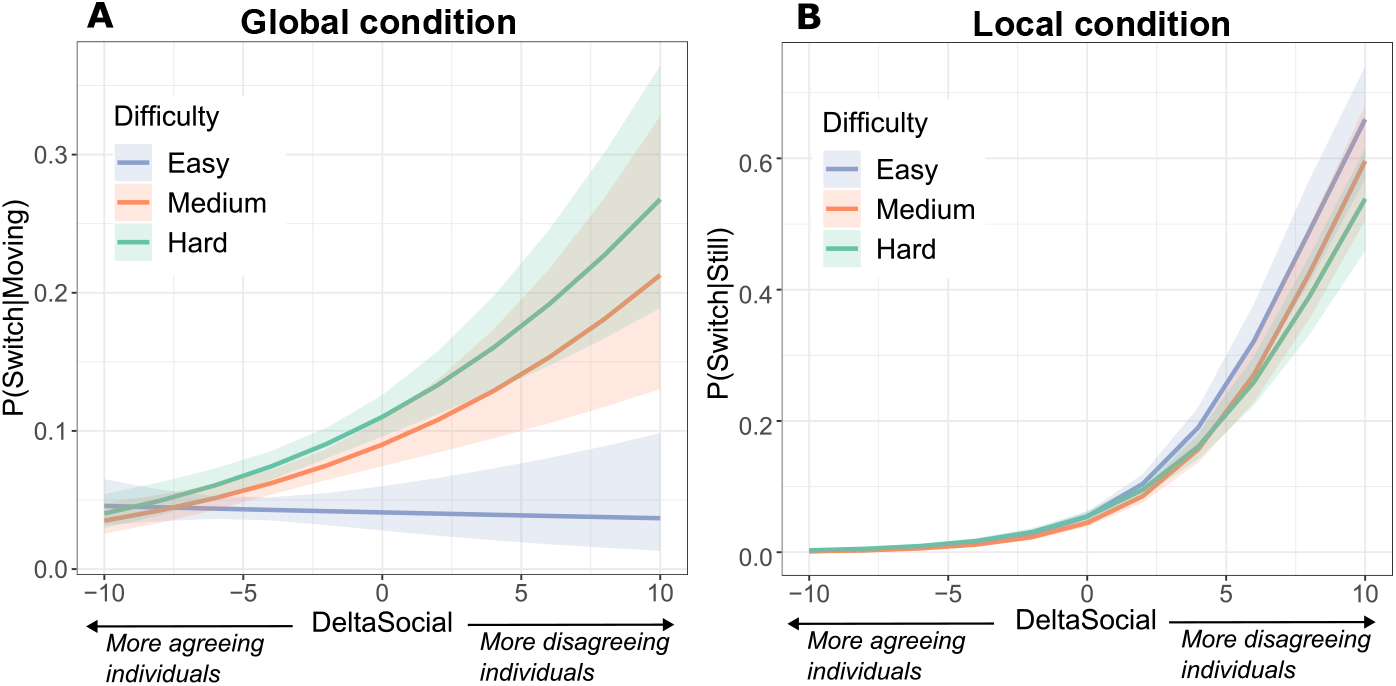
Likelihood to switch opinion under disagreeing social information. We assessed individuals that were moving towards or occupying a candidate site, for different difficulty levels. The likelihoods are estimates of the LMM. The maximum group size in the data was 10, and so the likelihoods at a `DeltaSocial` of -10 and 10 are extrapolations by the LMM. A) In the *global* condition, the probability of switching opinions while moving, according to the degree of disagreeing social information (`DeltaSocial`). B) In the *local* condition, the probability of switching opinions while occupying a candidate site, according to the degree of disagreeing social information (`DeltaSocial`).

In the *local* condition, participants often visited multiple options before converging on one. In this condition, the probability to move away from a currently occupied option (Fig. 3B) also increased as more individuals supported an alternative (*χ*^2^ = 2485.48, *p <* 0.0001). However, no significant interaction with difficulty level was found (*χ*^2^ = 3.89, *p* = 0.14), possibly because environmental information was less accessible, and thus it was harder for participants to estimate the difficulty of the task they were trying to solve. In short, the results indicate that participants’ choices were influenced by social information to a large extent, and the use of social information was modulated by the difficulty of perceiving the correct answer, but only if participants had a way to estimate that difficulty.

A model fit on data segments of the global information condition, across difficulty levels showed that, as participants moved closer to an option (see Fig. 4A), they also became less likely to switch opinions (main effect *β* = 0.01, SD = 0.001, *t* = 12.37, *p <* 0.0001). The effect of distance interacted with the degree of disagreeing social information (*β* = -5e-04, SD = 2e-04, *t* = -2.79, *p* = 0.005), indicating that the effect of distance was attenuated when social disagreement was higher. There was no significant interaction effect of distance with a quadratic term of the degree of disagreeing social information, suggesting that the attenuation was well represented by the linear relationship.

**Fig 4.**
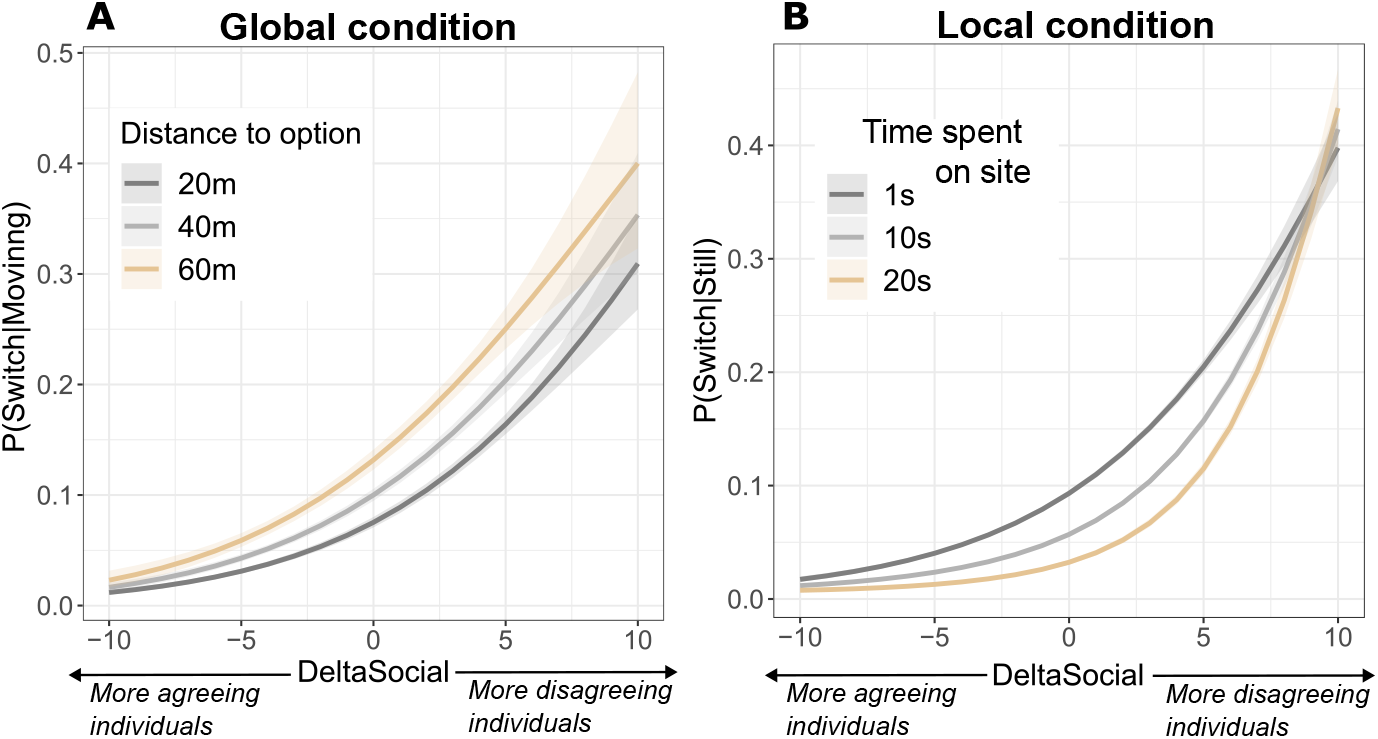
Dependence of switching opinion on spatiotemporal factors. A) The probability of switching opinion while moving, depending on the degree of disagreeing social information (`DeltaSocial`) and the distance to the currently selected site. B) The probability of switching opinion while occupying a candidate site, depending on the the degree of disagreeing social information (`DeltaSocial`) and the time already spent occupying that site.

In the local condition, When a participant was already occupying a candidate site (see Fig. 4B), the elapsed time spent there negatively affected the probability of switching opinions (*β* = - 0.05, SD = 0.0021, *t* = -24.77, *p <* 0.0001), indicating that participants were more likely to leave a site immediately after arriving than they were later on. There was also a positive interaction effect of time spent occupying a site with a quadratic term of `DeltaSocial` (*β* = 4e-04, SD = 1e-04, *t* = 6.11, *p <* 0.0001). This indicates that the effect of occupation time was attenuated by large positive or negative values of social disagreement. In short, social information use was importantly modulated by both spatial positions and sunk costs (i.e., reluctance to change after having spent time at a certain site).

### Behavior of informed individuals

The *informed minority* trials were more often successful and reached a consensus faster than trials without informed individuals (see “Reaching consensus on the empirically best option”). Informed individuals thus conveyed information to others through their movements. To investigate how they did so, we quantified differences in movement behavior between informed and uninformed individuals.

We quantified to what degree each participant either approached or moved away from the group’s center of mass (COM), as an indicator of (in)dependence from the group’s social influence. As expected, informed individuals were more likely to travel away from the COM than uninformed individuals, indicating that that they were more independent. Firstly, a positive model coefficient for distance to the COM (*β* = 0.13, SD = 0.02, *t* = 6.16, *p <* 0.0001) showed that individuals had on average a higher approach speed to the COM when they were further away from the COM, confirming that the COM had influence. Secondly, the distance to the COM, difficulty level (*easy, medium, hard*), and whether an individual was informed, all had significant main (*p* values *<* 0.0001), two-way interaction (*p* values *<* 0.0001), and three-way interaction (*p <* 0.03) effects on approach speed to the COM. Finally, the model coefficients showed a significant negative effect of being informed on the approach speed to the COM (*β* = -3.78, SD = 1.52, *t* = -2.49, *p* = 0.0127).

Informed individuals were also more likely than uninformed individuals to move directly to a candidate site without following a group majority, further confirming that informed individuals were more independent. This pattern was clearest in trials that were more difficult and in which uninformed individuals had less perceptual information. Firstly, positive model coefficients for the two-way interaction between distance to the COM and difficulty (*β*_medium_ = 0.18, SD = 0.03, *t* = 6.17 and *β*_hard_ = 0.14, SD = 0.03, *t* = -5.15, both *p* values *<* 0.0001) indicate that individuals were generally more influenced by the group in trials with higher difficulty. Secondly, negative coefficients for the three-way interaction effect between distance to the COM, difficulty, and whether an individual was informed (*β*_medium_ = 0.2, SD = 0.08, *t* = -2.57, and *β*_hard_ = 0.17, SD = 0.08, both *p* values *<* 0.031) indicate that the increased effect of distance to the COM with difficulty was smaller for informed individuals than uninformed individuals. Indeed, Fig. 5 shows that, for uninformed individuals but not for informed individuals, the approach speed to the COM depended more strongly on an individual’s distance to the COM when the difficulty level was higher.

**Fig 5.**
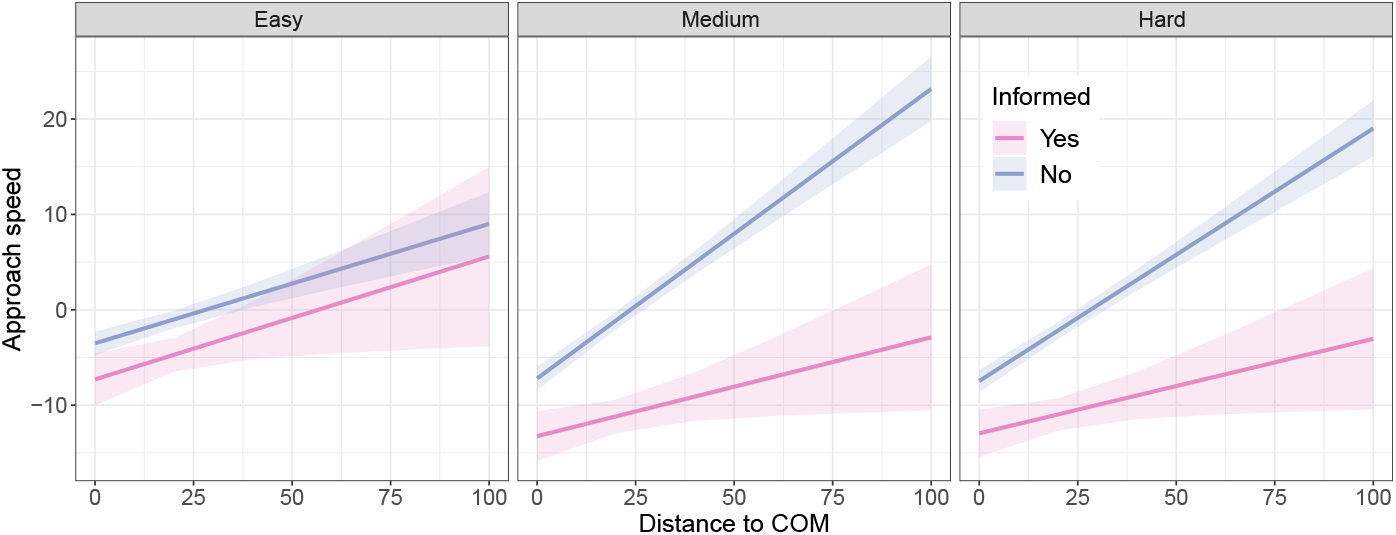
Effect of the group position on informed versus uninformed individuals. The difference in approach speed to the group center of mass (COM) according to the distance to the COM, for informed and uniformed individuals, in *informed minority* trials of different difficulty levels.

### Intentional persuasion

Due to the freedom of movement that participants had in our paradigm, we qualitatively observed many different behaviors. The anecdotal example presented in Fig. 6 shows how one informed individual engaged in an active persuasive behavior that consisted of leaving the correct site, approaching the incorrect majority on the site they were occupying, and then returning to the correct site. After initiation of the return movement, other participants gradually followed the informed individual to the correct site. Informed individuals presumably made such movements with the intention of persuasion (rather than because of being socially influenced themselves), as they were already informed of the correct option at the beginning of the trial. Four additional qualitative observations of persuasion behavior are presented in S4 Appendix.

**Fig 6.**
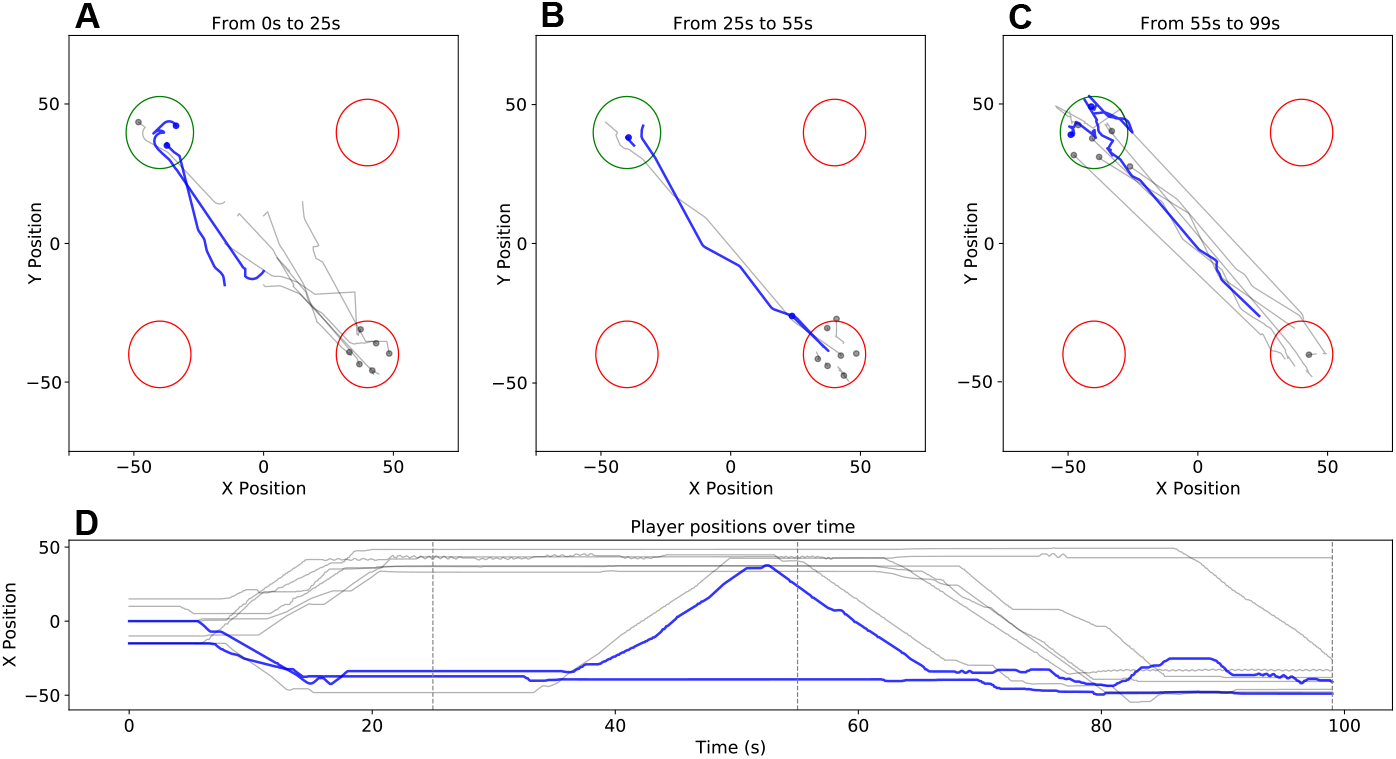
Example persuasive behavior by an informed individual. A–C) Trajectories during different phases in the trial (timestamps above each plot indicate trial phase). Trajectories of informed individuals are shown in blue, uninformed in gray. The correct site is shown in green, incorrect in red. D) The *x* position of each participant plotted over time. The vertical dotted gray lines indicate the end times of each of the three trial phases, corresponding to A–C.

## Discussion

In this study, we assessed whether groups of participants could reach a consensus on the empirically largest of four candidate sites without verbal communication and instead using movement-based interactions. In all conditions, the groups on average had an above chance performance for ending up on the largest site (i.e., *>*25% success rate, as there are 4 options). In most conditions, performance was well above chance (nearing or surpassing 75% success rate). The speed with which they chose an option reflected the difficulty of evaluating perceptual information about the site sizes (i.e. quality).

Perhaps surprisingly, trials with an incorrect consensus took on average a similar or longer time to reach a consensus than trials with a correct consensus; there was no speed–accuracy trade-off in our data. Speed–accuracy trade-offs have previously been observed in individual human decision making and in collective decision making by simple social animals and artificial agents [51–54]. It is not yet clear whether such a trade-off is present in some scenarios of collective human decision making, but so far, evidence for this is lacking. A previous study with movement-based group decisions in humans also did not find a speed–accuracy trade-off, but this was mostly because there were almost no occasions where groups landed on an incorrect site [14]. In our experiments, two factors might be related to the lack of an observed speed–accuracy trade-off. First, the longer time to reach a consensus for incorrect trials was possibly due to stubborn individuals who had high confidence in their (correct) answer and resisted conforming to the incorrect majority. Second, it is possible that the fastest decisions in our results were simply not fast enough to observe a decrease in accuracy. In house-hunting ants, it has been observed that the group will sometimes sacrifice accuracy to reach a faster decision when the environmental conditions are adverse, for example by lowering their quorum thresholds [52]. Future work could investigate whether incentivizing ultra-fast decisions in human groups could result in decreased accuracy, and therefore whether human groups might have a different trade-off distribution than social insect colonies (e.g., lower accuracy both for decisions that are too fast and too slow).

The presence of a small minority of informed individuals (2 individuals out of 7–10) aided a group in reaching the correct site quickly, even though the conditions were otherwise like those of the *global* trials, and therefore uninformed individuals could observe the full environment easily. Informed individuals also moved away from the group more often, indicating that their choices were more independent. This is reminiscent of behaviors by informed individuals previously observed in moving human groups, who spent less time following others than uninformed individuals [15, 16]. Also when informed individuals were present, the harder trials took longer to reach a consensus than the easier ones, indicating that informed individuals were not followed indiscriminately—indeed, uninformed individuals were not told whom was informed nor in which trials informed individuals were present. This is reminiscent of the behavior of poorly informed individuals observed in some site-selecting social insects. For example, poorly informed ants have been shown to still contribute to the colony’s decision, as the probability of an individual initiating recruitment to a site depended on the site’s quality even when the individual was poorly informed [40].

Both harder and easier trials were faster when informed individuals were present, and the independence of movements by informed individuals was more strongly displayed in harder trials, which supports previous findings that greater information discrepancy between informed and uninformed individuals can trigger leader–follower behaviors [55]. Although informedness was artificially induced in this study, it was not a simple all-or-none distinction, as uninformed individuals still made observations and could sample options differently based on their behaviors. In previous work, we have shown that such gradual differences in informedness between participants can be reflected in movement trajectories during collective decision making [33]. Still, it is important to note that the information discrepancies in this study are less complex than those that would likely be found in many real-life situations, which often involve multi-criteria decisions with undefined tradeoffs.

In fish shoals, potential leaders balance socially oriented behaviors (staying with the group) with goal-oriented behaviors (moving towards the candidate site) to drive the group towards correct consensus [56] in a way that could be seen as *laissez-faire* leadership rather than *active* leadership (e.g., active persuasion) [57]. In contrast, our results show some instances in which individuals attempted to actively persuade others: the persuader would closely approach the site being occupied by the other(s) and try to trigger them to leave. The process underlying this persuasion technique could be one of reciprocity (i.e., if I make the effort to move towards you, you might reciprocate by moving back with me). Indeed, the reciprocity and fairness norms that sometimes contribute to social influence are also applicable to physical effort [58–60]. In these cases, it is important that the persuader’s movement towards the recipient(s) is correctly interpreted by the recipient(s) as a persuasion attempt, rather than the persuader switching opinion to the site the recipient(s) are occupying. If the movement is misinterpreted, it might counterproductively reinforce the opinion the persuader is trying to dissuade (see S9 Fig in S5 Appendix). Further research could investigate the social inference capacities necessary to correctly interpret the intent with which such movements are made [61].

In our results, active persuasion attempts were also sometimes made using erratic movements as attempts to display confidence in opinion (see S7 Fig in S5 Appendix). This displayed confidence might result in a larger persuasive force for the advertised opinion, reminiscent of the relationship between waggle dance intensity and influence in honeybee decision making [3]. In honeybee communication via dances, changes in dancing are well-studied and both rate and duration have been shown to be important [3, 62]; studies have also investigated the mechanisms by which dancers increase their directional agreement as part of the site selection process [37]. Future research could focus on disentangling the relative importance and effectiveness of laissez-faire and active leadership in collective human decision making, and deciphering the mechanisms by which movement-based cues can be modulated, for more effective communication or to be more directionally aligned on the way to consensus. Movement-based leadership and influence can occur in many nuanced ways, even in our setup with limited communication abilities. Inferring influence from movement behavior can become even more complex in real-world situations, for example when inferring influence at different spatial and temporal scales [63]. Additionally, the correct decision in this study was based on empirical information, but the results might have been very different if consensus had needed to be reached about subjective preferences, in which case normative conformity largely drives social influence [64, 65]. Real-life situations can also involve a combination of empirical evidence and subjective preferences, as well as decision criteria that are complex or conflicting, resulting in unknown effects on the balancing of information and normative conformity.

In our experiments, the rate of opinion switching was lower in the *local* condition than in the *global* condition during movements, but higher when occupying a candidate site. This suggests that when information is readily available, initial decisions are made during initial movements, while, when information is less readily available, individuals explore before making use of social and individual information, using a kind of voting procedure by occupying preferred sites. Since participants did not have unlimited time to explore all options, they were presumably governed by an exploration–exploitation trade-off [66]: they could either invest time in exploring and therefore improving their individual information quality or move to an option selected by exploiting available social information (for example, by using the number of site occupants and the time spent there to infer site quality, as in [28]). Previous work has shown that groups can optimize their collective performance by adaptively balancing the use of individual exploration and social information, in a resource foraging task [67].

The probability of switching opinion while moving was significantly modulated by social influence (Fig. 3). This modulation was stronger in trials of higher difficulty, when perceptual information was more ambiguous (resembling previous findings that individuals are more likely to follow incorrect peers when the task is more difficult [26, 68]). However, in the *local* condition, we found no such interaction when an individual was occupying a candidate site rather than moving. This might have been caused by individuals being unaware of the task difficulty — and therefore immune to the effect of difficulty modulating social influence susceptibility — because they selected a candidate site without having explored the whole environment. Decision making was also modulated by site proximity: individuals were less likely to switch opinions when closer to the site they were currently moving towards. This reflects the “commitment effect” observed during embodied individual decision making (an internal positive feedback mechanism whereby people become more committed to a choice as they continue to approach it [29] and switching becomes more costly as the distance to the current opinion decreases and the distance to each alternative increases [69]), and suggests that the same effect is present during embodied group decision making. Multi-agent simulations have shown that such positive feedback mechanisms are conducive to reaching consensus in movement-based decisions [18]. We also found that participants were less likely to switch opinions after occupying a candidate site for a longer period of time, potentially indicating a sensitivity to sunk costs [31]. However, we observed this switching resistance (the effect of sunk costs) to be weaker in the presence of large opposing majorities.

In summary, we observed that reaching a consensus on empirical information when individuals have different opinions requires both positive feedback, to amplify an opinion to spread it through the group, and negative feedback, to trigger individuals to leave their opposing selection in favor of the majority opinion [70, 71], reminiscent of positive and negative feedback mechanisms in animal groups [18, 72]. For example, the attenuation of sunk-cost sensitivity by strong social influence could ensure disagreeing individuals do not block consensus. Indeed, when observing that your current commitment does not result in a consensus, it can be effective to reduce individual commitment in favor of social information. In our setup, participants’ use of social information was likely shaped by their goal to reach a consensus within a deadline, as individual concessions to majority opinions can depend on pro-social motivation [73] and the time pressure experienced by individuals [74]. It should be noted that our findings result from a very uniform sample consisting of WEIRD [75] university students. These results do not necessarily generalize to a broader population. Especially participants from cultures that differ on dimensions of individualism and power distance [76] might yield results that importantly differ from our WEIRD sample. Additionally, our sample of university students might have gaming experience, virtual interaction and navigation skills, or comfort with graphical user interfaces that might differ from other population samples and thus might have influenced the observed individual behaviors and group-level outcomes. Future work could investigate similarities and differences across cultures and different population subgroups, as well as across different analog and virtual task setups.

In conclusion, even though humans were not using any verbal communication in the present collective decision-making study, several different cognitive mechanisms still contributed to their modulation of social information. Although complex adaptive group behaviors might sometimes result from simple individual behaviors [16], explaining how human collective behavior emerges from individual behavior in more challenging settings requires taking into account more elaborate cognitive mechanisms [77]. Future research could focus, for example, on identifying the social inference mechanisms that contribute to adaptive collective decision making in embodied scenarios [28, 61, 78]. This study has begun an investigation into the interplay of individual exploration and social influence in human groups attempting to reach a consensus by pooling empirical evidence. However, we still understand far less about best-of-*n* decision making in humans than in several species of social insects, for which best-of-*n* decision making has been studied for decades in the context of nest-site selection. The results presented here indicate that the same macrophases of best-of-*n* decision making—individual exploration, information pooling, and reaching a consensus or quorum—are present in both humans and social insects. However, far more investigation would be required to determine the similarities to specific animal species, as even very similar species have been shown to have important differences (e.g., the red dwarf honeybee Apis florea versus the honeybee Apis mellifera [37]). Achieving an understanding of best-of-*n* decision making in humans equivalent to that of the most-studied insect species (Apis mellifera, Temnothorax albipennis) would require, at least, investigation of varying group sizes, varying the difficulty to discover the options, trade-offs between different aspects of quality, trade-offs between quality and the difficulty to discover the option, accuracy under ultra-fast or otherwise adverse conditions, and mechanisms that modulate the use of intentional cues. If future work addresses this gap, the resulting understanding of human group behavior could help improve a more general understanding of how species-common and species-specific aspects of best-of-*n* decision making are delineated across species, from humans to social insects.

## Materials and Methods

### Ethics statement

Participants were recruited from the student pool of Université Libre de Bruxelles and completed the experiment in return for course credits. Participants provided informed concent in written form on the experiment platform prior to starting the experiment. All experiment sessions were approved by the ethical committee of the Université libre de Bruxelles (permission 126/2020).The data collection was done in April and May 2021.

### Experiment setup

The experiments were conducted using HuGoS, our custom-developed platform for online experiments with multiple human participants in embodied scenarios inspired by swarm intelligence research [79]. The platform allows participants to connect remotely via their personal computer, joining a shared virtual environment by logging in at the same time. Participants interact with the platform similarly to an online multi-player game, using a keyboard and mouse and viewing the shared environment through the first-person views of their avatars. In this setup, participants were assigned to simple robot vehicle avatars that all looked alike; there were no visible differences between avatar bodies, such that participants would have been able to distinguish between them.

Each group of participants was scheduled to log in to the online platform at a specific time. Despite the aim of recruiting groups of 10 participants, connection problems and no-shows made that we had variable group sizes. In total, we had 9 sessions with 7 participants, 11 with 8, 8 with 9 and 4 with 10. Sessions with fewer than 7 participants were excluded from the analysis. Furthermore, inactive individuals (who did not control their avatar at all during an entire trial) were removed from the experiment for the subsequent trials. There were 263 participants in total (mean self-reported age = 20.6, SD = 4.1; self-reported as 215 female, 45 male, 3 other).

After logging into the game lobby, participants completed the mini Big 5 personality questionnaire [80] (not considered in the present analysis). The participants then watched a tutorial video explaining the task and then completed five practice trials. During the tutorial and practice trials, participants receive the following instructions: *In this task you have to find a treasure. The treasure is always hidden under the largest site. You can dig up the treasure if you are all on the same site. But watch out, the treasure is leaking! The faster you find the treasure, the more gold will be left*. If participants reached a consensus on the correct option, a treasure chest appeared and they accumulated *gold* points. The amount of *gold* points they received was 100 minus the time they took to complete the trial. If they took longer than 100 seconds, they saw the message *Too slow* and they did not receive any *gold* points. If they converged on the wrong option, they saw the message *The treasure was not here* and did not receive any *gold* points.

At the beginning of each trial, all participants started from one of several randomly assigned starting positions located within 20 meters from the environment’s center (see Fig. 1). In total, the environment size was 150 m by 150 m. The four options on which participants could possibly converge were placed at (−40 m,-40 m), (−40 m,40 m), (40 m,-40 m), and (40 m,40 m). Three of the options had a radius of 12 m. Depending on the difficulty of the trial, one of the four options had a radius of either 13 m (*hard*), 14 m (*medium*), or 15 m (*easy*). Participants controlled identical avatars with ground dimensions 3 m by 5 m that were shaped like bulldozers (Fig. 1). Participants could move forward or backward at a fixed speed of 8 m/s by pressing the ‘up’ or ‘down’ arrow keys, and they did not move when no key was pressed. They could rotate their avatar at *π* rad/s with the ‘left’ and ‘right’ arrow keys. See [79] for more details on avatar control.

## Analysis

### Statistics

For all analyses, we used either linear mixed models (LMMs) or generalized linear mixed models with a binomial response variable (GLMMs). We used the lme4 package in R to fit the models [81]. For continuous dependent variables such as time to reach a consensus and inter-player distance, we used LMMs. For binary dependent variables such as ‘successful’, we used GLMMs. For all group-level analyses, we included one datapoint per trial. The group ID and the number of participants in the trial were included as random effects to account for systematic behavioral differences between groups and between different group sizes. We also included the type of avatar that participants controlled as a random effect in the model. This factor was varied between groups but was discarded from the analysis.

For the individual opinion switch analysis, we included one datapoint per participant per second that they were either moving towards a candidate site or occupying a candidate site. The unique ID for that participant was included as a random effect for that participant.

Following each fitted GLMM, we conducted an analysis of variance (ANOVA) using Type II Wald *χ*^2^ tests to assess the significance of each fixed effect and its interactions within the models. We opted for a Type II analysis because it quantifies the distinct contribution of each model term, independent of the order in which they were entered into the model. This method evaluates the contribution of each model term, taking into account the presence of other terms but not their potential interactions. Consequently, it allows for the assessment of the independent significance of each fixed effect, untangled from its interactions with others.

Additionally, we conducted post-hoc tests using estimated marginal means (EMMs) of each GLMM using the `emmeans` package [82]. For each comparison between factor levels, we used the `pairs` function with the false discovery rate (fdr) adjustment to control for multiple comparisons. All post-hoc tests were two-sided.

### Descriptive results

Each trial could have three possible outcomes: 1) consensus on the correct opinion 2) consensus on the incorrect opinion, or 3) no consensus within the time limit. To determine the probability of success, we designated a trial as successful if it ended with a consensus on the correct opinion (1) and unsuccessful in the other cases (2 or 3). To assess differences between time to reach a consensus between conditions and difficulty levels, we only looked at the time to reach a consensus of successful trials.

### Position-based group measures

We assessed the average inter-player distance of participants as follows:

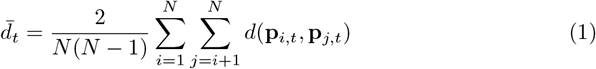

In the above equation, 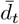 is the inter-player distance at time *t*, and *d*(**p**_*i,t*_, **p**_*j,t*_) is the Euclidean distance between participant *i* and participant *j* at time *t*.

Complementarily, we assessed the average distance of participants to the option that will eventually be chosen by the group:

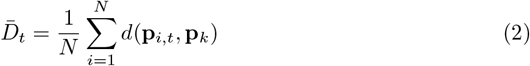

In the above equation, *D*_*t*_ is the average distance between the participants and the option at time *t*, and *d*(**p**_*i,t*_, **p**_*k*_) is the Euclidean distance at time *t* between participant *i* and the candidate site *k* that will eventually be chosen by the group.

### Quantifying opinions

Participants never explicitly indicated their support for one of the options. In order to facilitate the analysis of the opinion dynamics, we extracted participants’ opinions based on their positions and movements. We transformed the position and movements into a time series that represents which of the four options an individual is supporting over time (see S2 Fig in S2 Appendix). A participant was considered to support a candidate site if either occupying it or moving towards it. A participant was considered to be moving towards a candidate site if the avatar speed was nonzero and the angle between the velocity vector and a straight line from the avatar to the center of the option was less than 30 degrees. A participant was considered to be occupying a candidate site if not moving and positioned within the boundaries of the candidate site.

### Opinion switching analysis

We analyzed the probability that an individual would switch its opinion based on the state of the decision making process. We analyzed the probability of opinion switches during movements *P*(`Switch | Moving`) and when occupying a candidate site *P*(`Switch | Still`). For the analysis during movements, we divided each participant’s opinion time series into one-second bins. Each second during which the participant was moving towards a candidate site without switching its opinion was saved as a sample with `Switch` = 0. If during a subsequent second the individual switched to an alternative option and kept moving towards the alternative option for at least one second, then this was saved as a sample with *Switch = 1*. For the switching analysis when occupying a site, we saved a sample with `Switch` = 0 for every second in which an individual occupied a candidate site and saved a sample with Switch = 1 if the individual in a subsequent trial started moving away from the option towards an alternative option and continued doing so for at least one second.

#### Switching with condition and difficulty

For every sample, we saved whether participants were either moving towards or located on the correct or the incorrect option. First, we assessed how the probability of switching opinions was modulated by the experimental manipulations. Therefore, we fitted a GLMM with `Switch` as a dependent variable and `Condition, Difficulty` and `OnCorrect` as independent variables (including interaction effects). `OnCorrect` is a binary variable that is 1 if the participant is traveling towards or standing on the correct option, and 0 otherwise. Results of this analysis are plotted in see S5 Fig in S4 Appendix.

#### Switching with social influence and difficulty

To assess whether the probability of switching opinions differed with different degrees of social influence (Fig. 3 and Fig. 4 in the results), we defined a measure of the degree of disagreeing social information, `DeltaSocial`, as the difference between the largest number of individuals supporting any one of the unselected options and the number of individuals supporting the selected option. `DeltaSocial` is calculated as:

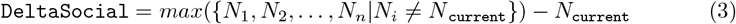

where *N*_*i*_ is the number of individuals supporting option *i* and `current` is the currently selected option. This number is positive when the number of disagreeing individuals (here taken as the ‘persuasive force’ for changing one’s opinion [83]) is larger than the number of agreeing individuals (i.e., the supportive force for one’s current opinion). We fitted a GLMM with `Switch` as a dependent variable and `OnCorrect, DeltaSocial`, and `Difficulty` as independent variables (including interaction effects). For opinion switches during movement, we only took into account trials from the *global* condition, since this was the condition where decision making is most likely to occur during movements. For opinion switches when occupying a candidate site, we only took into account trials from the *local* condition since this was the condition where the collective decision depended on participants switching between options.

#### Spatiotemporal modulations

In order to assess the degree to which spatiotemporal factors could modulate the participants decision-making process, we assessed to what degree the probability for switching opinions was influenced by a participant’s current position in space and the time it already spent at a candidate site (Fig 4 in the results). For opinion switches during movements, we assessed to what degree the probability of switching opinions was modulated by a participant’s distance to the option it was currently moving towards. We fitted a GLMM with `Switch` as a dependent variable and `Distance, DeltaSocial`, and `DeltaSocial`^2^ as independent variable (including interaction effects). We included a quadratic term of social influence in the model to capture the effect of the absolute majority size since both large positive values (when most participants agreed with the individual’s current opinion) and large negative values (when most disagreed) might similarly attenuate the effect of distance on the switching probability. For opinion switches when occupying a candidate site, we calculated the time spent at that site so far and fitted a GLMM with `Switch` as a dependent variable and `TimePassed, DeltaSocial`, and `DeltaSocial`^2^ as independent variables (including interaction effects).

### Movement behavior for informed and uninformed individuals

To evaluate differences in behavior between informed and uninformed individuals (Fig. 5), we quantified to what degree they approached or went away from the group’s center of mass (COM). Moving away from the COM can be interpreted as independent movement, while movements toward the COM could be seen as socially influenced movements. The group’s center of mass at time *t* is given by:

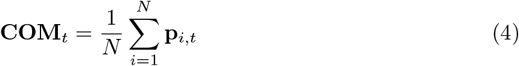

We quantified the approach speed of a participant within a certain time window as the negative derivative of the participant’s distance to the COM. This derivative was multiplied by the norm of the participant’s absolute velocity, in order to only capture the participants’ active movements towards the COM, rather than movements of the COM relative to the player:

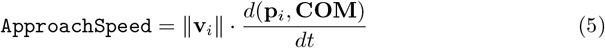

where ‖ **v**_**i**_ ‖ represents the absolute speed of the participant relative to the environment, and *d*(**p**_*i*_, **COM**) represents the distance between the participant and the group COM.

## Supporting information

**S1 Fig. Proportion of correct, incorrect, and uncompleted trials**. Reported in S1 Appendix.

**S2 Fig. Example of how trajectories are transformed into opinion time series, in a trial with 7 participants**. Reported in S2 Appendix.

**S3 Fig. Group cohesion during trials that ended with a correct consensus**. Reported in S2 Appendix.

**S4 Fig. Completion times for correct and incorrect trials**. Reported in S3 Appendix.

**S5 Fig. Probability of opinion switching for different difficulty levels and conditions**. Reported in S4 Appendix.

**S6 Fig. Example of majority influence during movement, in the** *global* **condition**. Reported in S5 Appendix.

**S7 Fig. Example of a “signaling” behavior**. Reported in S5 Appendix.

**S8 Fig. Example of fast consensus, in the** *informed minority* **condition**. Reported in S5 Appendix.

**S9 Fig. Example of a failed persuasive behavior by an informed individual.c**Reported in S5 Appendix.

**S1 Appendix. Correct, incorrect, and uncompleted trials**.

**S2 Appendix. Group cohesion on the way to consensus**.

**S3 Appendix. Completion times for correct and incorrect trials**.

**S4 Appendix. Individual resistance to change of mind**.

**S5 Appendix. Anecdotal examples of qualitative behaviors**.

