## Supplementary information for "Best-of-*n* decision making by human groups"

#### S1 Appendix: Correct, incorrect, and uncompleted trials

Fig. S1 shows the number of trials that have ended in a correct consensus, an incorrect consensus, or which have not resulted in a consensus by the time limit.

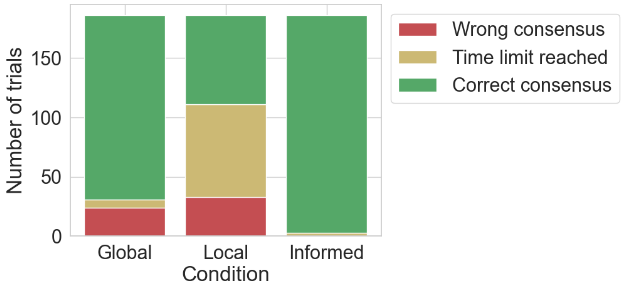


**Figure S1: Proportion of correct, incorrect, and uncompleted trials.** The proportion of trials of each condition that have ended in a wrong consensus (red), a correct consensus (green) or that have not converged before the time limit (yellow).

### S2 Appendix: Group cohesion on the way to consensus

In the successful trials, we analyzed to what degree participants tend to favor similar options throughout the trial. For this analysis, we transformed the movement trajectories of participants into a discrete opinion time series based on the candidate sites they were either standing on or moving towards (see Methods; Fig. S2) and we quantified the degree to which participants adopted the same discrete opinion throughout the trials. We also calculated the average inter-individual distance between participants' avatars as a measure of group cohesion (Fig. S3).

The average size of the majority (i.e., the largest subgroup of individuals that agree on one option) during the trial ($\chi^2(2) = 8.61, p = 0.01$) as well as the average inter-individual distance ($\chi^2(2) = 23.359, p < 0.0001$) during the trial were modulated by the difficulty level of the trial, suggesting that participants' balancing of individual exploration and social information use was modulated by the their level of confidence in their empirical observations.

Both majority size ($\chi^2(2) = 350, p < 0.0001$) and inter-individual distance ($\chi^2(2) = 1473, p < 0.0001$) were also modulated by the conditions. The average majority size was smaller ($t = -12.8, p < 0.0001$) and the average inter-individual distance was larger ($t = 32.18, p < 0.0001$) in the *local* condition than in the *global* condition. This shows that participants engaged in more exploration and information gathering in the *local* condition, suggesting a resistance to early use of social information when collecting empirical evidence requires exploration.

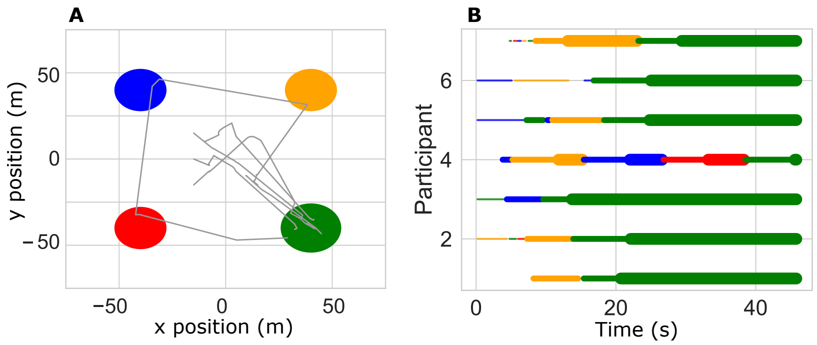


**Figure S2: Example of how trajectories are transformed into opinion time series, in a trial with 7 participants.** A) Trajectories traveled by all participants during the example trial. B) The color of the line indicates which candidate site each participant supports during one trial. Thin lines indicate that a participant is looking towards that candidate site, medium thick lines indicate participants are moving towards it, and thick lines indicate participants are positioned on top of it.

The inter-individual distance was marginally smaller in the *informed minority* condition than in the *global* condition ($t = -2.03, p = 0.04$), suggesting that informed individuals can to some degree steer the decision process in a way that maintains group cohesion. Moreover, visual inspection of Fig. S3 shows that the most pronounced decrease in inter-individual distance in the *informed minority* condition occurred in the *hard* difficulty level, suggesting that participants were more likely to be influenced by their peers when having lower confidence in their own empirical observations.

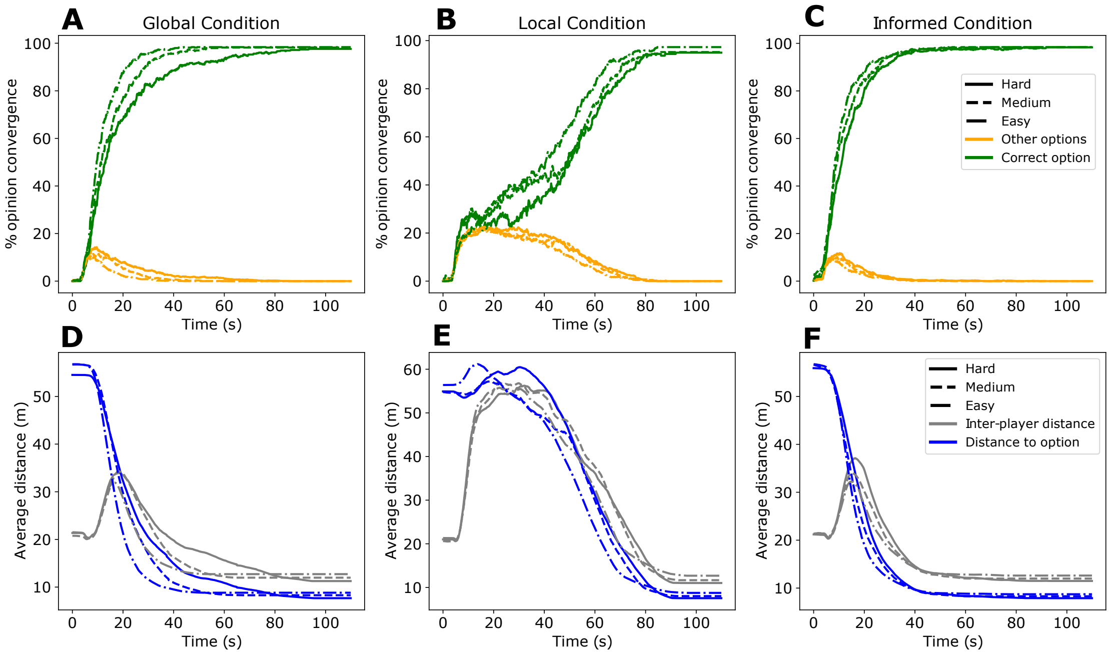


**Figure S3: Group cohesion during trials that ended with a correct consensus.** A-C) Average opinion convergence over the course of a trial. The green lines indicate the growing support for the correct candidate site. The yellow lines indicate the average support for any of the three other options. The different line styles indicate averages of trials with different difficulty levels. D-F) The blue lines indicate the average participant distance to the correct candidate site. The gray lines indicate the average inter-player distance between participants throughout the trials. The different line styles indicate averages of trials with different difficulty levels.

### S3 Appendix: Completion times for correct and incorrect trials

Fig. S4 shows the completion times of correct and incorrect trials for the different difficulty levels in the *local* and *global* conditions. Data from the *informed minority* condition is not shown since there were no incorrectly completed trials.

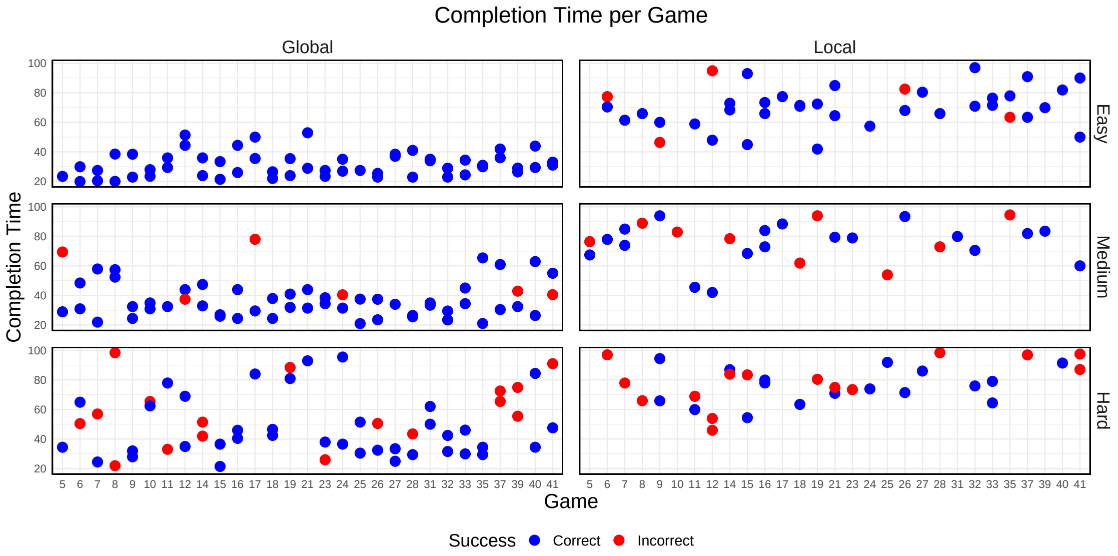


**Figure S4: Completion times for correct and incorrect trials.** Each colored dot represents the completion time of one trial. Different subplots indicate different conditions and difficulty levels. The color of the dots indicates whether a trial was completed with a correct consensus (blue) or an incorrect consensus (red). The x-axis represents the different experimental sessions. In each experimental session, the group completed two trials of each condition and difficulty level. Games with less than two data points per subplot had surpassed the time limit in the missing trials.

### S4 Appendix: Individual resistance to change of mind

To assess how individual decision-making behavior is impacted by the different experimental conditions, we quantified how easily participants switch from the option they currently support to an alternative option. To do so, we subdivided the opinion time series (see Fig. S2) of each participant into one-second epochs, and calculated the probability that a participant would change their current opinion within one epoch given the experimental conditions and current state of the experiment. We first analyzed how the individual probabilities for switching opinions varied between conditions and difficulty levels without taking into account social influence strength (see Fig. S5). The GLMM fit showed that switching probability was meaningfully modulated by the condition, difficulty level, whether participants were currently moving towards the correct candidate site, and all the interaction effects (type II ANOVA, all p values $< 0.0001$). Participants were more likely to switch opinions when moving toward the incorrect site than when moving toward the correct site ($z = 24.597, p < 0.0001$), indicating that, on average, participants could distinguish between the correct and incorrect sites, whether by using their own direct observations or by observing the actions of others.

Participants in the *local* condition were less likely to switch away from movements towards incorrect opinions than in the *global* condition ($z = -4.07, p < 0.0001$), possibly reflecting the fact that participants in the *local* condition were not able to observe the empirical quality of the site before reaching it. Conversely, the probability of switching while occupying a candidate site was higher in the *local* condition than in the *global* condition ($z = -12.9612, p < 0.001$), suggesting that participants in the *global* condition were more confident in their choice and therefore more resistant to changing their opinion than in the *local* condition.

Participants were more likely in the *informed minority* condition than in the *global* condition to switch away from an incorrect option, both while moving towards it ($z = 4.07, p < 0.0001$) and while occupying it ($z = 6.50, p < 0.0001$), suggesting that informed individuals had greater social influence than uninformed individuals, on average reducing their peers' resistance to change of mind.

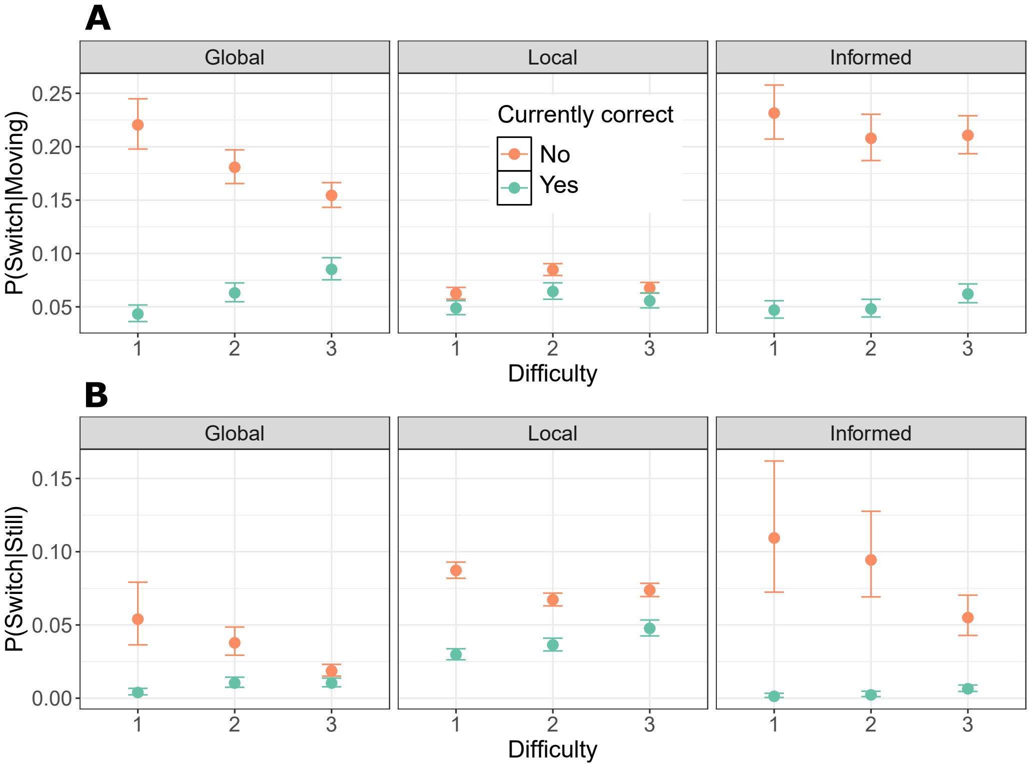


**Figure S5: Probability of opinion switching for different difficulty levels and conditions.** A) The probability of switching opinions in a given second while moving towards a candidate site. B) The probability of switching opinions in a given second when occupying a candidate site.

### S5 Appendix: Anecdotal examples of qualitative behaviors

We here show some anecdotal examples of participant behavior. Fig. S6 shows a common pattern of collective behavior in the *global* condition: individuals initially travel to different options, but disagreeing individuals eventually change their movement direction to be in line with the majority opinion. Fig. S7 shows an example of a "signaling" behavior that was often observed in the *local* condition. When individuals were confident in the option they were currently standing on, they made circular movements, as can be seen by the circle in the trajectory in Fig. S7B and the oscillation in Fig. S7D. Fig. S8 shows a pattern that was commonly present in the *informed minority* condition. The informed individuals initiate movement towards the correct option at the early stages of the trial and are then followed by the uninformed individuals. Fig. S9 shows a situation that was only rarely observed. An informed individual moves towards individuals on the incorrect option (supposedly in an attempt to convince them to switch options; see main paper) but is then followed by others who also leave the option that the informed individual initially supported (possibly due to misinterpreting the persuasive behavior of the informed individual as an opinion switch). The informed individual is then not able to convince the uninformed individual to change options before the end of the trial.

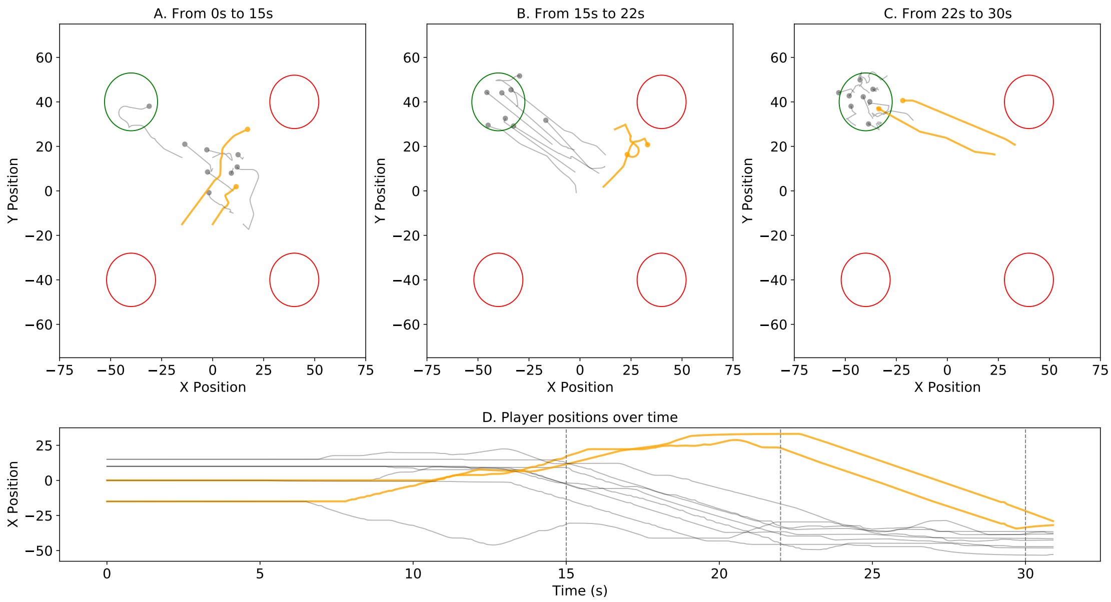


**Figure S6: Example of majority influence during movement, in the *global* condition.** A–C) Trajectories of different phases in the trial. Timestamps above each plot indicate the trajectory phases. The trajectories of two individuals that initially disagreed with the majority are shown in orange. The trajectories of the other participants are in gray. The correct option is indicated with a green circle; the incorrect options are indicated in red. D) The $x$ position of each participant is plotted over time. The dotted gray lines indicate the part of the trials that are plotted in A–C.

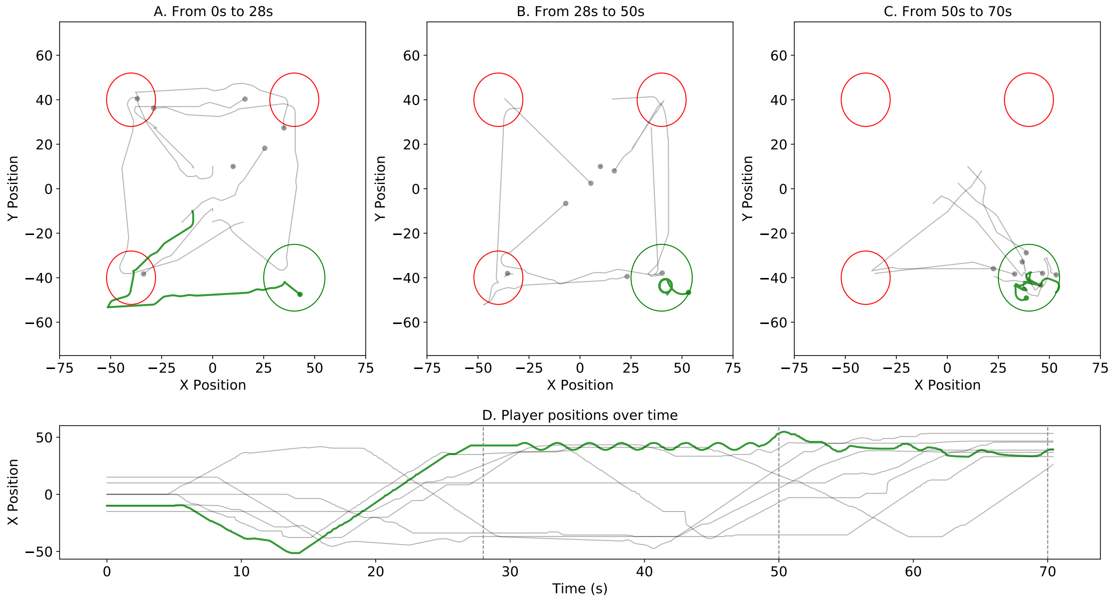


**Figure S7: Example of a "signaling" behavior.** A–C) Trajectories of different phases in the trial. Timestamps above each plot indicate the trajectory phases. Trajectories of the individual that performs circular "signaling" movements are shown in green. The trajectories of the other participants are in gray. The correct option is indicated with a green circle; the incorrect options are indicated in red. D) The $x$ position of each participant is plotted over time. The dotted gray lines indicate the part of the trials that are plotted in A–C.

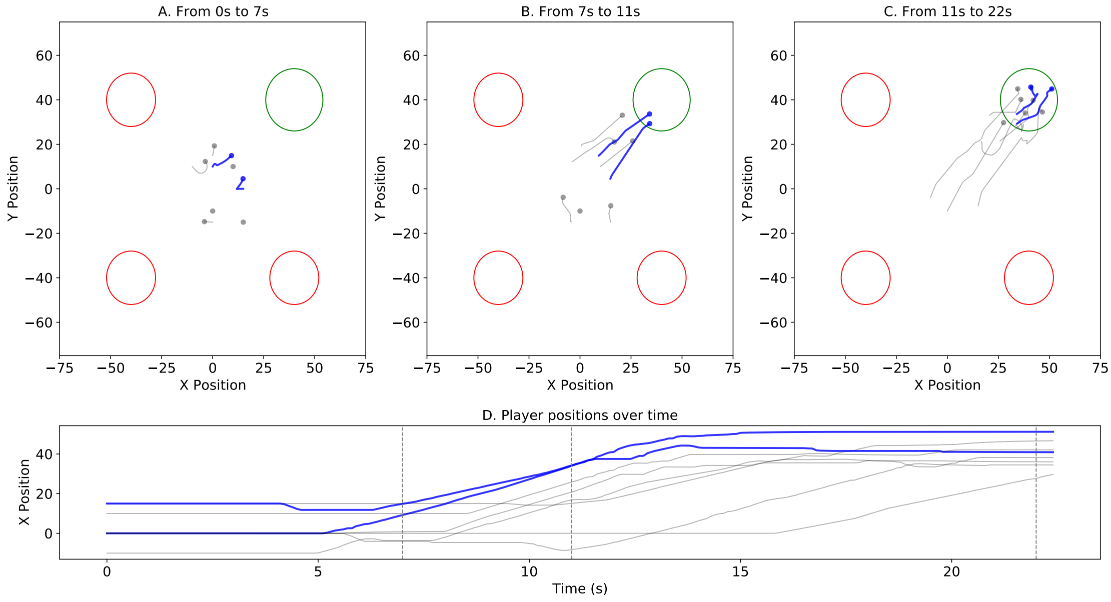


**Figure S8: Example of fast consensus, in the *informed minority* condition.** A–C) Trajectories of different phases in the trial. Timestamps above each plot indicate the trajectory phases. Trajectories of informed individuals are shown in blue. The trajectories of the other participants are in gray. The correct option is indicated with a green circle; the incorrect options are indicated in red. D) The $x$ position of each participant is plotted over time. The dotted gray lines indicate the part of the trials that are plotted in figures A–C.

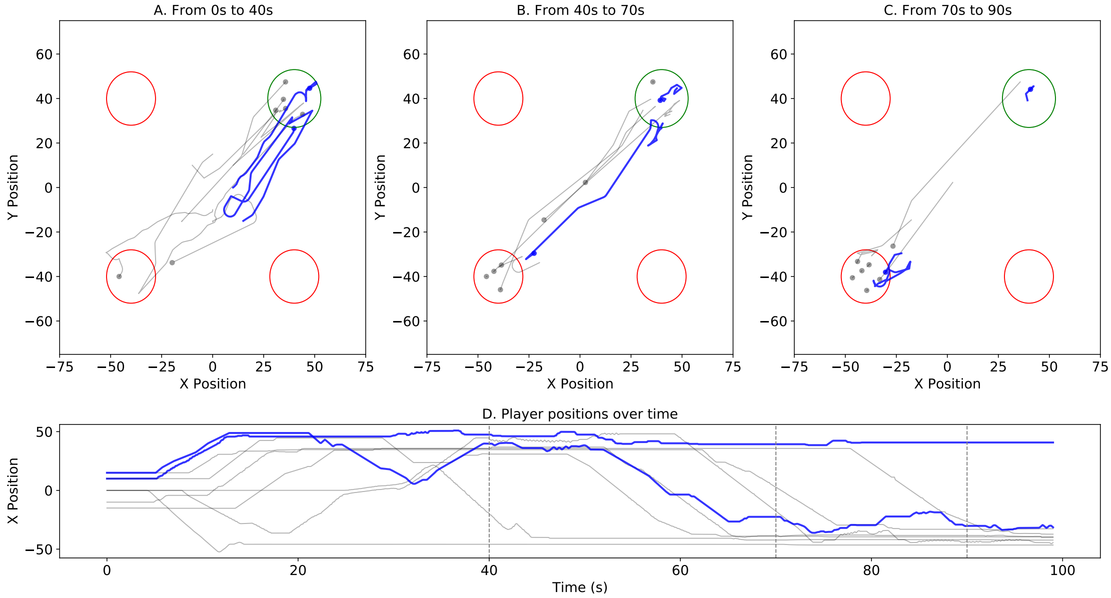


**Figure S9: Example of a failed persuasive behavior by an informed individual.** A–C) Trajectories of different phases in the trial. Timestamps above each plot indicate the trajectory phases. Trajectories of the two informed individuals are shown in blue. The trajectories of the other participants are in gray. The correct option is indicated with a green circle; the incorrect options are indicated in red. D) The $x$ position of each participant is plotted over time. The dotted gray lines indicate the part of the trials that are plotted in A–C.
